# TracktorLive: an integrated real-time object tracking and response system

**DOI:** 10.64898/2026.03.12.711471

**Authors:** Pranav Minasandra, Vivek H. Sridhar, Dominique G. Roche, Isaac Planas-Sitjà

## Abstract

Real-time tracking and automated response systems are essential for standardising experiments, reducing observer bias, and improving reproducibility in studies of movement and behaviour. However, existing solutions face significant challenges: AI-based tracking systems require expensive hardware and impose computational delays, creating challenges for closed-loop experiments; existing real-time tracking tools lack standardised implementations for response delivery; and steep learning curves limit accessibility for users without programming or computer vision expertise. Here, we introduce TracktorLive, an open-source Python package designed to overcome these challenges through concurrency and a modular, ‘cassette’-based architecture. TracktorLive leverages traditional computer vision techniques to perform image-based object detection without the need for expensive hardware or deep learning. By parallelizing object tracking and response delivery into separate, concurrent server and client processes, the software minimizes frame processing time, enabling rapid, real-time analysis and response delivery. User-friendly ‘cassettes’—portable code snippets that can be copy-pasted into scripts—enable users with minimal programming experience to implement complex workflows for use in experiments and practical applications. We demonstrate TracktorLive’s utility through several applications, including microcontroller-based stimulus delivery for location-dependent manipulations; conditional video recording that activates only during events of interest; kinematic-based response triggering using real-time velocity computations; and multi-cassette experimental designs combining multiple functionalities. Detailed tutorials are provided to familiarize users with TracktorLive’s operation and functionality, and a growing library of cassettes supports diverse applications for processing both real-time and pre-recorded video. We validated the software by comparing its response timing to human experimenters in a stimulus delivery task involving two fish species, where TracktorLive demonstrated consistently higher accuracy and lower variability, particularly for fast-moving subjects. Beyond experimental biology, TracktorLive’s unique architecture and versatility could support many different applications in fields ranging from neuroscience to wildlife management. As an open-source software combining accessibility, modularity, and computational efficiency, TracktorLive can help democratize real-time tracking and automated response systems across disciplines.

## 1. Introduction

Data collection during studies involving movement and behaviour often relies on human intervention and direct observation, which introduces bias (Box 1), can lead to experimenter-driven disturbances [1–3], and requires a significant investment of time and effort on the part of the experimenter. A common approach to increasing measurement accuracy and reducing bias is to record subjects on video and implement *post hoc* tracking using manual or automated solutions [4–6]. However, many experiments also necessitate dynamic interactivity between the experimenter, the arena, and the subjects. For instance, a shrew may need to be released into a maze at a specific time, and a reward dispensed if it attempts to solve the maze [7]. Similarly, a stimulus might need to be delivered when a school of fish enters a specific area of the experimental arena [8,9]. In more complex setups, a continuous display might need to be updated dynamically so an insect perceives simulated movement in a virtual landscape [10,11]. Such experiments will benefit from real-time tracking and automated responses [12,13], which offer a promising approach to standardizing experiments, reducing observer bias, and improving experimental efficiency and reproducibility.

In the last fifteen years, major advances have been made in tracking the positions of cells [14,15] and animals [16–22] from video data. These tracking tools have helped overcome the constraints of traditional observational methods and resulted in a bloom of proprietary and open-source solutions in fields ranging from experimental biology to pest management [16,23–26].

Recent advances in automated, image-based tracking have resulted in the development of a host of tracking techniques, both with and without Artificial Intelligence (AI). However, tools are lacking that allow experimenters to design a response system that can decide on and execute real-time responses based on the tracked locations of objects in video. When researchers have had to adopt automated response delivery systems, they have typically developed a specific solution—such as the use of infrared beams or RFID tags— that may not be applicable to other experimental setups [12,27,28]. Most recently, methods focussing specifically on real-time animal tracking have become available, which rely on deep learning-based approaches [21,29,30]. While highly accurate and robust, these solutions are computationally intensive and may impose a recurrent lag for processing each frame. As a result, AI-based tracking has improved *post hoc* analyses. However, numerous challenges need to be overcome for their widespread applicability in closed-loop experiments, where stimuli must be delivered immediately in response to location or behavioural changes, particularly when conducted on lower-end hardware. Some real-time tracking software can interface with external hardware, such as Raspberry Pi or Arduino boards, to facilitate stimulus delivery [9,26,29], but these systems are often designed for specific experimental setups and lack standardised implementations for closed-loop experiments, limiting their generalisability, accessibility, and broader adoption.

Key challenges standing in the way of a generalisable, real-time tracking and response software are (i) the computational overhead incurred by AI-based programs for consecutive tracking and response operations at each frame, both of which can be time-consuming steps requiring expensive hardware; and (ii) the steep learning curve that must be mastered by researchers, often with no experience in computer vision, who must learn to use existing libraries and program actions in a way that harmonises with computer-vision-based tracking algorithms.

Here, we introduce TracktorLive, an open-source python package that uses concurrency (the ability to execute more than one program or task simultaneously) and a modular, ‘cassette’-based approach, to overcome these challenges. We elaborate on TracktorLive’s architecture and use (Section 2); detail several examples of applications (Section 3); discuss its strengths, limitations, and future prospects (Sections 4 and 5); and compare its performance with that of human experimenters (Supplementary Information).

### Box 1.

**Reducing observer bias in experimental studies involving animal movement and behaviour**

Minimizing observer bias has been a long-standing challenge for studies involving animal movement and behaviour—in ecology, medicine, neuroscience, and experimental biology, among other fields [31–33]. In particular, data collection during behavioural experiments often relies on direct observation, which is prone to bias and experimenter-driven disturbances [1–3]. A common approach for reducing these effects is video recording experiments to avoid the presence of an observer, and using ‘blind’ protocols, in which the experimenter is unaware of a subject’s identity and treatment group [2,34–36].

Despite the benefits of blind protocols [32,33,37–39], blinding is often impractical—for example, when a single researcher runs an experiment, or in field conditions where time and resources to conduct experiments are limited. Similarly, the presence of an experimenter near test animals is sometimes unavoidable, particularly when a stimulus must be presented or triggered—for instance, when deploying a decoy predator to elicit an escape response [8,40], or for delivering food as positive reinforcement [41]. Even when a stimulus is operated remotely, experimenters may still introduce bias due to individual variability in training and experience, their subjective perceptions and interpretations of animal behaviours, and variability in reaction times when triggering stimuli. Additionally, subtle gender biases have been documented among experimenters, leading to different behavioural outcomes in test subjects [42,43]. Given these challenges, real-time tracking and automated stimulus delivery offer a promising approach to standardizing experiments and reducing observer bias. See Supplementary Information for an experimental comparison of human-mediated and automated response delivery using an early version of TracktorLive. The results show a clear benefit of using automated response delivery, which reduced the experimental error within and among trials (Fig. S3).

## 2. Software

### 2.1. How TracktorLive works (software architecture)

TracktorLive is available for installation as a one-command pip-installable python package (`pip install tracktorlive`). As a fully-fledged python package, the capabilities of TracktorLive are vast. To fully leverage this growing range of capabilities, TracktorLive is designed with a command-line focus, avoiding users and developers being restricted by operating-system specific norms in graphics. TracktorLive is primarily designed for GNU/Linux operating systems like Ubuntu or Pop! OS, but the software, the examples, and the ‘cassettes’ (see below) have all been installed and tested on Windows 11 (using the Windows Subsystem for Linux, WSL). Detailed installation instructions and relevant notes for all operating systems are available on the GitHub repository.

The utility of TracktorLive is derived from three important design choices to overcome key challenges of implementing real-time tracking and response software.

First, at its core, TracktorLive runs Tracktor [16], an image-based object detection and tracking freeware that does not rely on deep-learning. Instead, Tracktor adopts traditional computer vision techniques including adaptive thresholding (to segment the object of interest from the background) and the Hungarian algorithm (to link individual identities between frames), making it relatively lightweight and eliminating the need for expensive hardware or specialised user training. Thanks to Tracktor’s minimalistic design, TracktorLive can easily be expanded with additional pre-processing steps such as masking and background subtraction to improve tracking performance.

Second, TracktorLive parallelises object tracking and actions through a semaphore-protected shared memory architecture, which substantially reduces the time taken for processing each incoming video frame (Fig. 1). ‘Semaphore’ locks prevent multiple parallel processes from simultaneously trying to write or read data from the shared memory (in what are called ‘race’ conditions), ensuring that only one process has access to tracking data at any time. The primary parallel process, a ‘Tracktor Server’, handles all tasks related to object tracking in incoming video frames, while separate parallel processes (‘Tracktor Clients’) are initiated to adjudicate response delivery. As a result of this parallelisation, real-time tracking and response is possible without incurring significant computational delay between consecutive frames.

**Figure 1:**
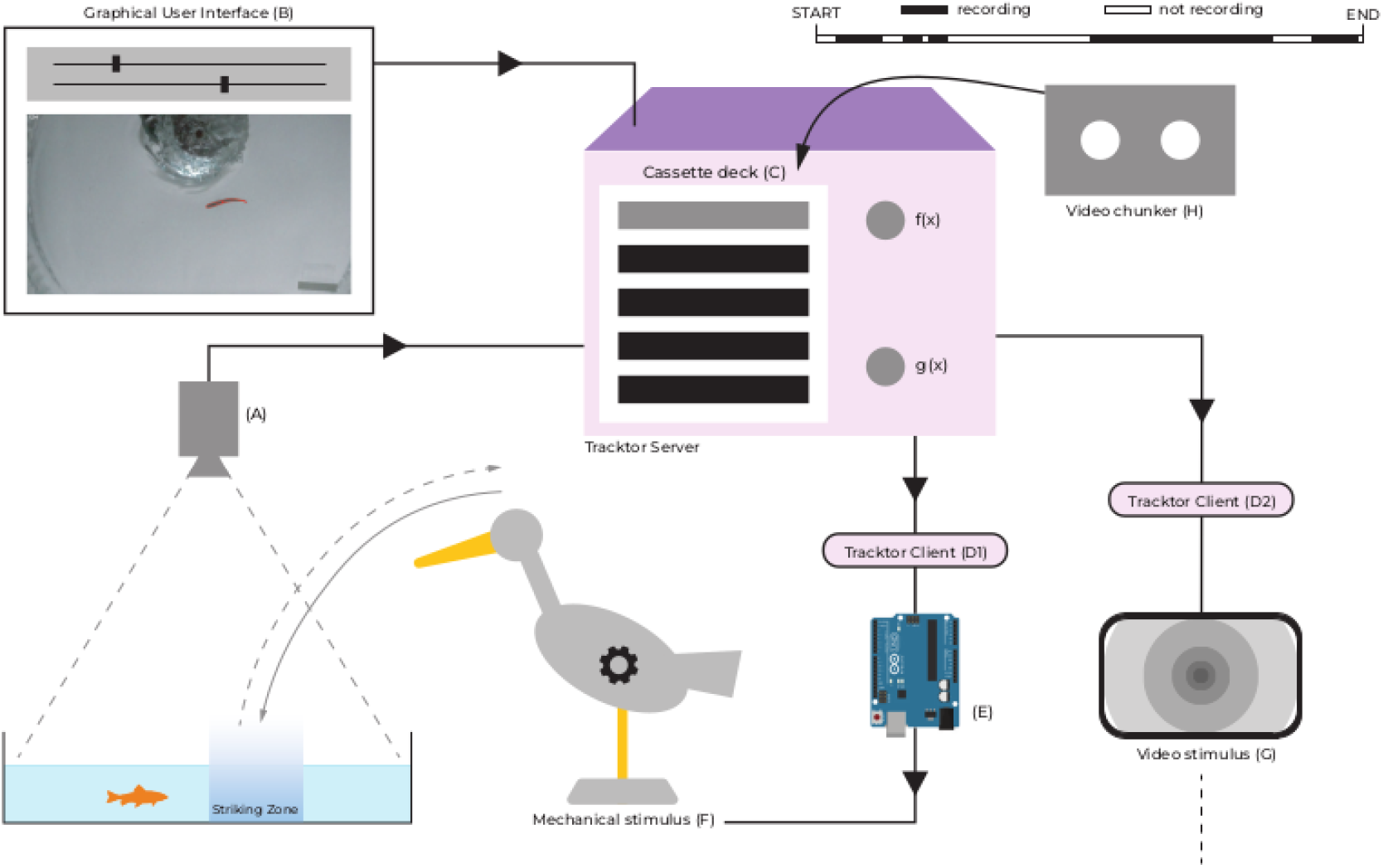
An overview of the TracktorLive pipeline. Frames from a live feed **(A)** recording subjects (here, a fish in a tank) are passed to a Tracktor Server, for which tracking parameters can be specified through a custom GUI **(B)**. The Tracktor Server’s behaviour is specified by the addition of ‘cassette functions’ to its on-board cassette deck **(C)**. The tracked location of each animal is shared with multiple Tracktor Clients **(D1, D2)**, which can drive response delivery. For example, client **D1** can send messages to an Arduino board **(E)**, which triggers a mechanical stimulus **(F)** to elicit an escape response from the fish when it is in a certain location. Meanwhile, Client **D2** can play a video **(G)** to trigger a looming stimulus when, for instance, the fish moves below a certain velocity for a specified duration. Sometimes, users may want to toggle a switch to save video dynamically in response to the animals’ positions. The Video Chunker cassette **(H)** can toggle whether video is saved to disk in response to the fish’s location, preserving disk space by saving video only when it is of interest. The figure illustrates only a small subset of the types of experiments TracktorLive can facilitate.

Third, TracktorLive is readily accessible to those unfamiliar with computer vision or advanced programming. The software has been designed to work with ‘cassettes’. Cassettes are small, portable snippets of code designed to be ‘dropped in’ scripts to achieve particular tasks (Fig. 2), such as video processing and response delivery. TracktorLive includes a command-line utility that automates simple tasks and circumvents the need for writing code for basic applications of the software (Fig. 2). The software also comes with a graphical user interface (GUI) that facilitates tuning the object detection and segmentation parameters prior to tracking. These features make implementing real-time tracking and responses accessible to users with minimal experience with the python programming language. A detailed description of TracktorLive’s architecture and design is provided as Supplementary Information and in Fig. S1.

**Figure 2:**
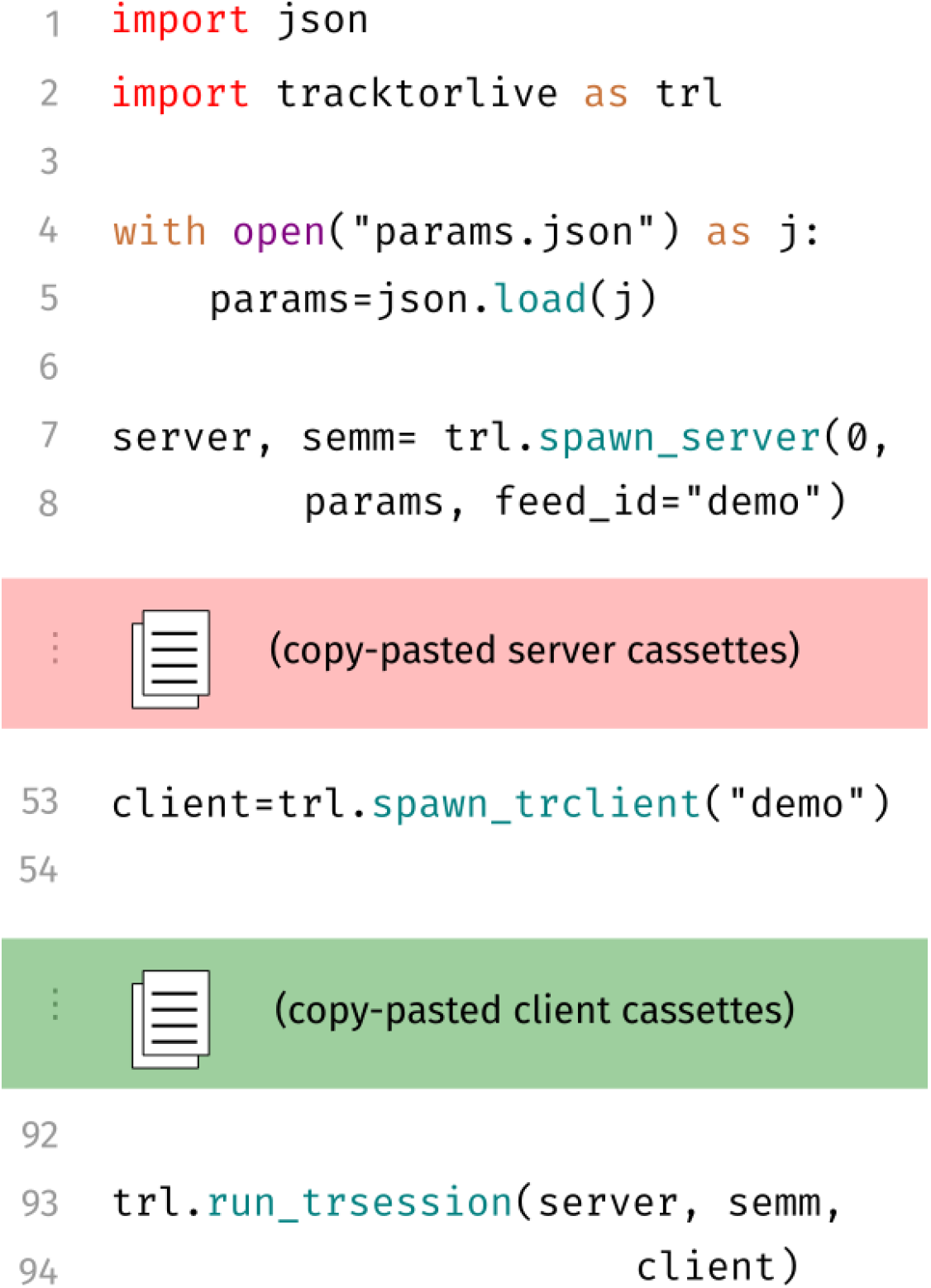
The standard structure of a TracktorLive script. Every TracktorLive python script starts with importing `tracktorlive` as `trl` (by convention) (L2). Then, after reading tracking-related parameters (L5), the Tracktor Server is declared (L7-8). Along with the server, a semaphore manager (`semm`) is automatically spawned. The user then customises the behaviour of the server by inserting server cassettes. These could, for instance, control when video storage is turned on, or edit the incoming video stream on-the-fly to improve the tracking performance (for example, adjust the brightness and contrast). Then, optionally, one or more clients are declared (L53). As with the server, the behaviour of the client is customised by adding client cassettes, which can handle visualisation, interfacing with other devices, or running commands in response to the movements of tracked animals. Finally, the server, semaphore manager, and client are all initialised together at the end of the script (L93-94). See Fig S2 for a visualisation of an actual script with complete code.

### 2.2. Running TracktorLive

The standard way of running TracktorLive is to write python scripts (Figs. 2 and S2) that declare a Tracktor Server (which does all the object tracking in parallel, behind the scenes) and, optionally, a number of Tracktor Clients, which handle response delivery. ‘Cassettes’ are modular code snippets that can be copy-pasted into scripts and stacked in any user-defined order to change the behaviour of the program to achieve specific goals. We currently provide over 30 cassettes for a broad range of applications (Table 1).

**Table 1:** TracktorLive Cassettes. These cassettes are available on the TracktorLive GitHub repository’s [Library of Cassettes](https://github.com/pminasandra/tracktorlive/tree/master/Library_Of_Casettes), and can be copy-pasted into existing scripts to achieve the desired behaviours (explained in the ‘Description’ column). User-specifiable parameters are displayed at the beginning of the code for each cassette to enable customization.

| Cassette | Description |
| --- | --- |
| Display Feed | Display current tracking from the server in real-time. Press `q` or `Esc` to close running display at any time |
| Add Circular Mask | Mask everything except a circle of specified position and radius |
| Add Custom Mask | Add a custom mask to every frame based on user-provided image file |
| Add Rectangular Mask | Mask everything except a rectangle of specified vertices |
| Also Preserve Audio | Extract audio from input video and muxes it into the output video at the end |
| Also Record Audio | Record audio with `ffmpeg` in a separate process and muxes it into the recorded video at the end |
| Background Subtract | Replace background pixels with white using an averaged background image |
| Boost Contrast | Brightness and contrast control |
| Channel Select | Preserve only one of the BGR channels |
| Color Select | Keep only pixels close to a user-defined BGR colour; mask out everything else |
| Delayed Start | Delay the start of tracking by a fixed amount of time |
| Display Final Vel Plot | Display and save a plot of velocities when we reach the end of the video file we are analysing |
| Display With Circ HI | Display current tracking from the server in real-time. Highlight a chosen circular region. Press `q` or `Esc` to close running display at any time |
| Display With Rect HI | Display current tracking from the server in real-time. Highlight a chosen rectangular region. Press `q` or `Esc` to close running display at any time |
| Dynamic Masking | Only look for individuals in vicinity of previous locations |
| Extract Background | Compute per-pixel variability to estimate a background image |
| Extract Specified Frames | Save as `jpg` all frames at specified indices |
| First Person Views | For each tracked individual, crop a fixed-size square that is rotated so the animal's heading points "up". Uses EMA + rate limiting to smooth orientation. Writes one MP4 per individual |
| Gaussian Blur | Apply a Gaussian blur to each frame before tracking |
| Message Arduino | When a user-defined function returns a character, transmit that character to a connected Arduino |
| Overlay Text From DataFrame | Overlay user-provided frame-wise text on video |
| Preserve Frame | Put non-edited frames in `framesbuffer` (useful for `dumpvideo` methods) |
| Print Directions | Calculate and print the direction based on position changes |
| Print Position | Print current position of the target individual |
| Record When Together | Record video only when two animals are within a specified distance of each other |
| Run Command On Condition | Run a shell command when a function returns True, respecting a time based cooldown rule |
| Time Lapse | Build a time-lapse by subsampling frames into a custom buffer and writing once at the end |
| Timesync | Create a parquet file of `time.time()` clock, useful for ms level sync |
| Update Plot | Display (and save) a plot every few seconds. Useful to monitor data in real time |

The behaviours of both the Tracktor Server and Tracktor Clients are fully programmable by inserting cassettes into the script. Experienced users and developers can easily expand existing cassettes and create new ones to accomplish additional tasks. In the spirit of open-source software, we explicitly welcome cassette contributions from users to be added to the Library of Cassettes. Cassette creators are specified and credited at the start of the code for each cassette. We expect TracktorLive’s capabilities to increase further as the community begins to adopt it and more cassettes are contributed.

## 3. Applications

Applications of TracktorLive’s real-time capabilities are vast, ranging from stimulus delivery when objects are in a given location or perform certain behaviours; recording and tracking movements only under specific conditions; live plotting data as an experiment unfolds; and activating mechanisms to open doors or deliver food; to implementing conditional workflows to alert experimenters by email or phone (Table 1).

Below, we briefly detail four examples of common applications in experimental scenarios (Fig. 3). Python scripts for these examples are available on the TracktorLive GitHub repository.

**Figure 3:**
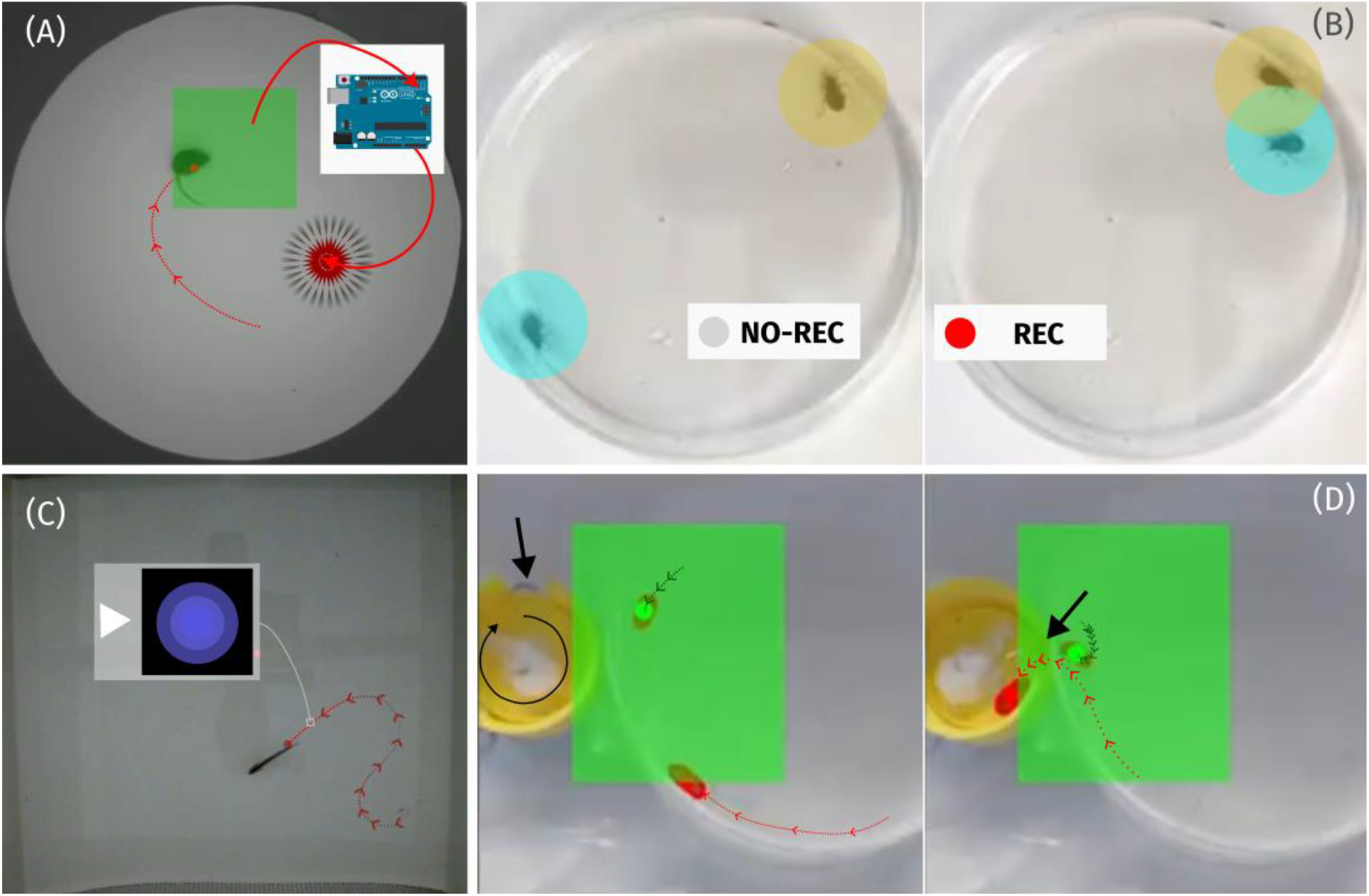
Examples of four applications of TracktorLive. **(A)** a threatening stimulus (represented here by the flashing of an LED) is triggered through an Arduino board when TracktorLive detects a mouse entering a specified area of interest; **(B)** recording to disk is turned on only when two leafbugs are within a threshold distance of each other; **(C)** a video that contains a looming stimulus is played when a fish exceeds a certain velocity; and; **(D)** a door (thick black arrow) is opened by driving a servo motor when two pillbugs simultaneously enter a specified area of interest.

### Example 3.1. Real-time experimental manipulations with microcontrollers

Microcontroller boards, such as Arduino boards (www.arduino.cc), are frequently used to design response systems that manipulate setups during experiments. TracktorLive can readily be programmed with a single cassette to communicate with a connected Arduino board. By programming the Arduino board to respond to incoming information from TracktorLive, real-time, automated experimental manipulations become feasible. Fig. 3A and Video S1 illustrate an example of an “agonistic stimulus” triggered in response to the location of an animal in an arena.

### Example 3.2. Conditional video recording based on event triggers

When experimenters are interested in specific, infrequent events, video recordings can become arbitrarily long. This can lead to large video files and a high storage capacity demand, as well as a heavy time investment on the experimenter’s part to visualise the entire recording and analyse specific events. In such scenarios, the Tracktor Server can be set up with a cassette to only record videos when events of interest occur. Fig. 3B shows an example where video storage is only triggered when two animals are in close proximity, and therefore likely interacting. In this specific example, the leafbugs only interacted for 2 min 12 sec within a 50 min period (see Video S2).

### Example 3.3. Stimulus delivery based on real-time computation of kinematics

Since TracktorLive computes real-time position, it is possible to perform quick computations on tracks using a cassette. In Fig. 3C, we compute the velocity of a fish and trigger the presentation of a looming stimulus when the fish exceeds a certain velocity threshold (Video S3). Note that the cassette used (‘Run Command on Condition’) could easily be programmed to respond to any kinematic condition (position, acceleration, orientation, and so on) and to trigger any other action; for example, sending the experimenter an email if the focal animal has been stationary for an unexpectedly long period of time.

### Example 3.4. Stacking cassettes for complex experiments

Finally, to illustrate the modular nature and stackability of TracktorLive cassettes, we combine examples 1 and 2 in a simple experiment where a door in the experimental arena is opened only when two pillbugs are in proximity to it (Fig 3D, Video S4). This example uses the compute proximity cassette highlighted in example 2 with an Arduino board operating the door via a servo motor, as in example 1.

Beyond the examples above, additional tutorials are available on the TracktorLive GitHub repository, demonstrating several other applications of the software (Table 2). A first set of tutorials explains the use of the command-line tool to perform quick tasks such as recording a video and tracking individuals under simple conditions. Subsequent tutorials demonstrate how to optimize tracking by adding cassettes for the adjustment of contrast and brightness and the application of masks. Finally, the last set of tutorials show how users, including those with minimal coding experience, can customize existing cassettes and create new ones on their own or with the help of large language models (LLMs). One example provided illustrates how LLMs can help users create cassettes to accomplish a difficult task: converting a recording of multiple individuals in an arena into bird’s-eye chase videos centred around each individual. This example demonstrates how a hypothetical user could readily solve a new, custom task like creating viewpoints that facilitate subsequent analyses such as pose estimation and behavioural classification [44]. This straightforward applicability of LLMs in the use of TracktorLive is facilitated by the modular and compact nature of cassettes. We also provide a ‘customGPT’ in the OpenAI framework on the TracktorLive GitHub repository, which we have tested with some cassettes in our Library of Cassettes. We specifically label cassettes that were created using this tool so users can compare them with cassettes we wrote ourselves. We emphasise that our inclusion of this tool is not an endorsement of the company or its LLMs, and code generated by this tool should be thoroughly tested (as with any LLM use). Finally, while TracktorLive’s novelty lies in its ability to perform real-time object tracking and response delivery, the software and all its cassettes are equally applicable to pre-recorded video.

**Table 2:**
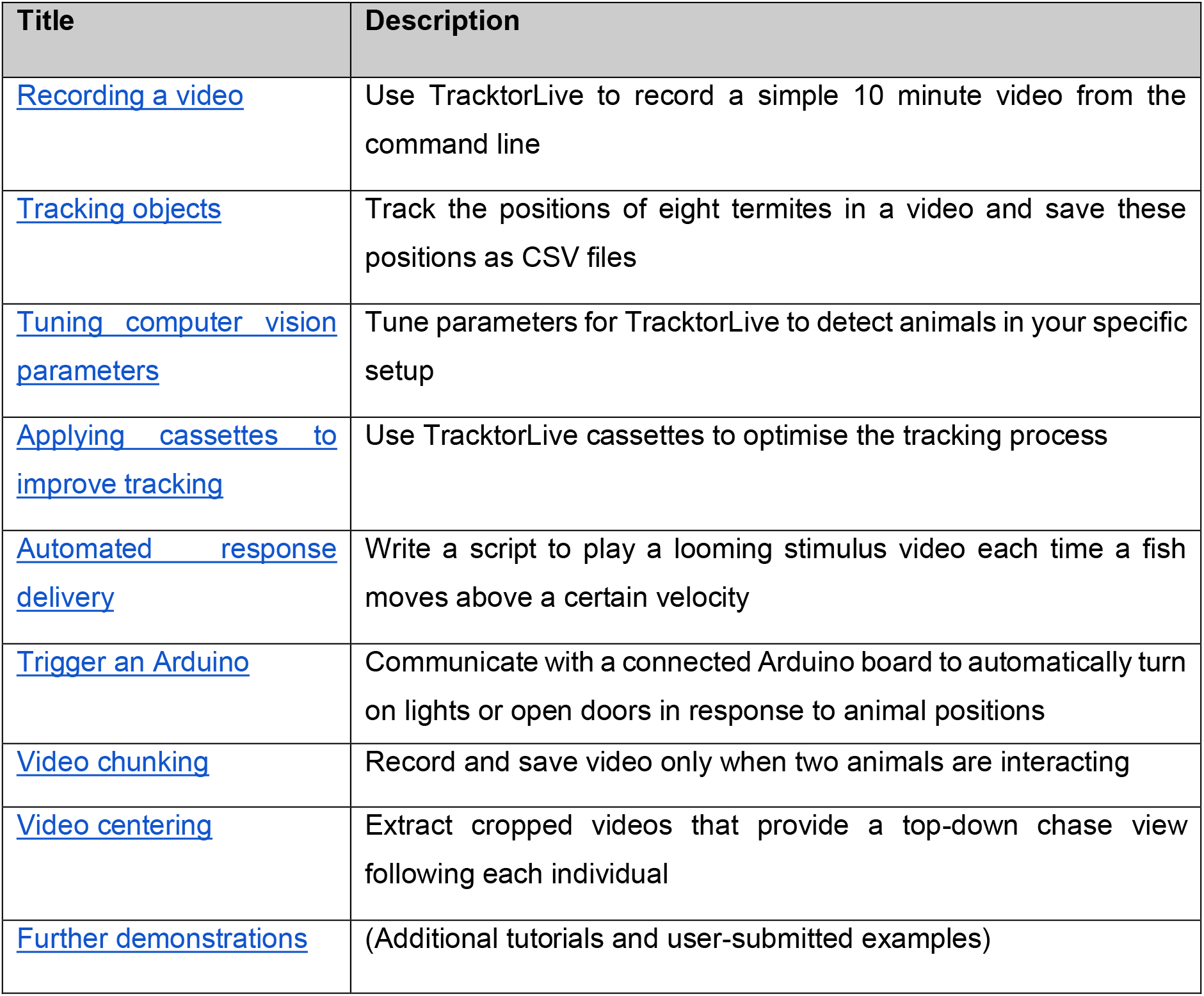
TracktorLive Tutorials. Tutorials are available on the TracktorLive GitHub repository to help users familiarize themselves with the software.

We encourage users to submit new applications of TracktorLive for inclusion as ‘Further Examples’ in the tutorials and to contribute to the “living” Library of Cassettes.

## 4. Limitations

TracktorLive uses Tracktor [16] under the hood, and therefore shares both its strengths and limitations. Tracktor can reliably track single individuals under varying experimental conditions, including in situations with uneven lighting and/or where the background is not static. It can also simultaneously track multiple individuals in the same or in distinct arenas. However, Tracktor—and by extension, TracktorLive—is not designed to solve crosses, occlusions and/or a change in the number of individuals in the arena: situations where multiple animals walk, fly or swim above or below each other, enter and leave the arena, or are occluded from the camera (for example, when entering a shelter or exiting the camera’s field of view). This is because Tracktor maintains the individual identities of unmarked objects using the Hungarian algorithm, a highly reliable technique provided there are no crosses or occlusions. In scenarios where crosses or occlusions can occur or the number of individuals is dynamic, TracktorLive can be used reliably only when tracking and response delivery does not depend on the identities of the individuals tracked.

In scenarios where maintaining individual identities is essential, the use of a traditional (non-AI) tracking algorithm becomes infeasible. However, the implementation of Tracktor within TracktorLive occurs within a single method (`Server._eachframe`) in the Tracktor Server object class, and an experienced developer could easily replace this core tracking method with a different, AI-based method (for instance, based on the “You Only Look Once” (YOLO) framework [45]). Given the modularity of TracktorLive, this modification would maintain the integrity and benefits of TracktorLive’s architecture as well as the use of the library of cassettes and command-line utility. Similarly, developers could modify TracktorLive’s tracking method to detect ArUco markers—open-source simplified QR codes that can be mounted on animals [25]—allowing to maintain individual identities without the use of AI when crosses or occlusions occur, or when the number of individuals in the arena changes.

Finally, while the stackability of cassettes is a strength of TracktorLive, stacking too many cassettes risks affecting the processing speed of the software. If users create and stack custom cassettes that are computationally demanding, they should actively monitor the software to ensure it behaves as expected, the main risk being that frames are skipped during processing.

## 5. Future prospects

Numerous potential applications of TracktorLive exist that we have not yet explored. Specifically, the integration of TracktorLive with low-cost embedded systems (e.g., Arduino), could enable a variety of closed-loop systems (setups where a biological process is monitored in real-time, and information is used to immediately adjust a treatment or stimulus). In organismal biology, these enhanced setups could enable better experimental design in the study of escape responses, animal cognition, or decision-making. Applications also extend across a wide range of domains beyond the study of animal movement and behaviour, from cell biology to ecological monitoring. For instance, closed-loop systems leveraging TracktorLive could facilitate experiments in which neural or cellular activity of labelled cells is linked to state changes (e.g. fluorescence) or movement, allowing real-time modification of environmental conditions based on detected signals [46]. Integrating TracktorLive with deep learning tools could also help support wildlife surveys and monitoring through automated recording and analysis of occurrences and activity.

TracktorLive can also be helpful to automate and optimize lab protocols. Using cassettes, a researcher can be notified when an experiment ends and schedule automatically triggered analyses scripts or the secure transfer of data and results to a remote location, digital repository or website. TracktorLive could also help improve animal welfare; for example, by automatically detecting sick/dead animals based on kinematics, or automatically regulating oxygen levels and water replacement, or providing environmental enrichment. Finally, enthusiasts could develop offline, privacy-friendly home applications for day-to-day use, or for educational purposes to promote scientific interest. The possibilities are vast (see Table S2).

## 6. Conclusion

TracktorLive addresses a critical gap in experimental biology by providing a low-cost, readily accessible solution for real-time tracking and responses to movement and behaviour. The modular architecture and efficient parallelised processing can support a diversity of experimental designs without expensive hardware or advanced programming skills. As the research community adopts TracktorLive, we anticipate the Library of Cassettes will continue to expand through user contributions that extend the software’s capabilities, including into new domains. The integration of TracktorLive with emerging technologies including, advanced imaging techniques, embedded systems and machine learning holds particular promise for developing innovative applications. Ultimately, by lowering the financial and technical barriers to real-time tracking and response systems, TracktorLive provides a scalable, accessible, and reproducible solution for studying movement and behaviour across diverse systems.

## Data availability statement

The data and the analysis script to reproduce our results are available on https://github.com/pminasandra/tracktorlive and the Open Science Framework repository [47], and were shared with the editors and reviewers upon submission.

## Ethics statement

The experiment adhered to ethical regulations of Tokyo Metropolitan University (Japan), which do not require ethical approval for observational studies using fish. Throughout the study, no animals were harmed or subjected to stressors other than being transferred from one aquarium to another. The principal investigator (I.P.S.) completed the animal training course at Tokyo Metropolitan University and ensured that the study was conducted following ethical guidelines aimed at minimizing any potential impact on the fish.

## Acknowledgements

P.M. and V.H.S. acknowledge support from the Max Planck Society (Max-Planck-Gesellschaft). V.H.S. additionally thanks the Department of Biology, University of Washington for their support. This study was also funded by the Japan Society for the Promotion of Science (JSPS KAKENHI 18F18806) awarded to I. P.-S.

## References

1. Marsh DM, Hanlon TJ. Seeing What We Want to See: Confirmation Bias in Animal Behavior Research. Ethology. 2007;113: 1089–1098. doi:10.1111/j.1439-0310.2007.01406.x

2. Tuyttens FAM, de Graaf S, Heerkens JLT, Jacobs L, Nalon E, Ott S, et al. Observer bias in animal behaviour research: can we believe what we score, if we score what we believe? Anim Behav. 2014;90: 273–280. doi:10.1016/j.anbehav.2014.02.007

3. Keaney TA, Jones TM, Mulder RA. An undergraduate classroom experiment illustrates an effect of observer bias on data collection in animal behaviour. Anim Behav. 2024;212: 175–182. doi:10.1016/j.anbehav.2024.03.013

4. Davidson JD, Sosna MMG, Twomey CR, Sridhar VH, Leblanc SP, Couzin ID. Collective detection based on visual information in animal groups. J R Soc Interface. 2021;18: 20210142. doi:10.1098/rsif.2021.0142

5. Gelblum A, Pinkoviezky I, Fonio E, Ghosh A, Gov N, Feinerman O. Ant groups optimally amplify the effect of transiently informed individuals. Nat Commun. 2015;6. doi:10.1038/ncomms8729

6. Sampaio E, Sridhar VH, Francisco FA, Nagy M, Sacchi A, Strandburg-Peshkin A, et al. Multidimensional social influence drives leadership and composition-dependent success in octopus–fish hunting groups. Nat Ecol Evol. 2024;8: 2072–2084. doi:10.1038/s41559-024-02525-2

7. Baldoni C. From summer growth to winter decline : brainsize, captive effect, and cognitive outcomes in the common shrew during Dehnel’s phenomenon. 2024 [cited 6 Mar 2026]. Available: https://kops.uni-konstanz.de/handle/123456789/72832

8. Herbert-Read JE, Ward AJW, Sumpter DJT, Mann RP. Escape path complexity and its context dependency in Pacific blue-eyes (Pseudomugil signifer). J Exp Biol. 2017;220: 2076–2081. doi:10.1242/jeb.154534

9. Planas-Sitjà I, Ioannou CC. State-behaviour feedbacks between boldness and food intake shape escape responses in fish (Gasterosteus aculeatus). Commun Biol. 2025;8: 227. doi:10.1038/s42003-025-07669-w

10. Sayin S, Couzin-Fuchs E, Petelski I, Günzel Y, Salahshour M, Lee C-Y, et al. The behavioral mechanisms governing collective motion in swarming locusts. Science. 2025;387: 995–1000. doi:10.1126/science.adq7832

11. Sridhar VH, Li L, Gorbonos D, Nagy M, Schell BR, Sorochkin T, et al. The geometry of decision-making in individuals and collectives. Proc Natl Acad Sci. 2021;118. doi:10.1073/pnas.2102157118

12. Humby T, Laird FM, Davies W, Wilkinson LS. Visuospatial attentional functioning in mice: interactions between cholinergic manipulations and genotype. Eur J Neurosci. 1999;11: 2813–2823. doi:10.1046/j.1460-9568.1999.00701.x

13. Bathellier B, Tee SP, Hrovat C, Rumpel S. A multiplicative reinforcement learning model capturing learning dynamics and interindividual variability in mice. Proc Natl Acad Sci. 2013;110: 19950–19955. doi:10.1073/pnas.1312125110

14. Piccinini F, Kiss A, Horvath P. CellTracker (not only) for dummies. Bioinformatics. 2016;32: 955–957. doi:10.1093/bioinformatics/btv686

15. Aragaki H, Ogoh K, Kondo Y, Aoki K. LIM Tracker: a software package for cell tracking and analysis with advanced interactivity. Sci Rep. 2022;12: 2702. doi:10.1038/s41598-022-06269-6

16. Sridhar VH, Roche DG, Gingins S. Tracktor: Image-based automated tracking of animal movement and behaviour. Methods Ecol Evol. 2019;10: 815–820. doi:10.1111/2041-210X.13166

17. Pérez-Escudero A, Vicente-Page J, Hinz RC, Arganda S, de Polavieja GG. idTracker: tracking individuals in a group by automatic identification of unmarked animals. Nat Methods. 2014;11: 743–748. doi:10.1038/nmeth.2994

18. Yamanaka O, Takeuchi R. UMATracker: an intuitive image-based tracking platform. J Exp Biol. 2018;221: jeb182469. doi:10.1242/jeb.182469

19. Torney CJ, Dobson AP, Borner F, Lloyd-Jones DJ, Moyer D, Maliti HT, et al. Assessing Rotation-Invariant Feature Classification for Automated Wildebeest Population Counts. PLOS ONE. 2016;11: e0156342. doi:10.1371/journal.pone.0156342

20. Chiara V, Kim S-Y. AnimalTA: A highly flexible and easy-to-use program for tracking and analysing animal movement in different environments. Methods Ecol Evol. 2023;14: 1699–1707. doi:10.1111/2041-210X.14115

21. Diwan T, Anirudh G, Tembhurne JV. Object detection using YOLO: challenges, architectural successors, datasets and applications. Multimed Tools Appl. 2023;82: 9243–9275. doi:10.1007/s11042-022-13644-y

22. Bartashevich P, Herbert-Read JE, Hansen MJ, Dhellemmes F, Domenici P, Krause J, et al. Collective anti-predator escape manoeuvres through optimal attack and avoidance strategies. Commun Biol. 2024;7: 1586. doi:10.1038/s42003-024-07267-2

23. Mathis A, Mamidanna P, Cury KM, Abe T, Murthy VN, Mathis MW, et al. DeepLabCut: markerless pose estimation of user-defined body parts with deep learning. Nat Neurosci. 2018;21: 1281–1289. doi:10.1038/s41593-018-0209-y

24. Herrera C, Williams M, Encarnação J, Roura-Pascual N, Faulhaber B, Jurado-Rivera JA, et al. Automated detection of the yellow-legged hornet (Vespa velutina) using an optical sensor with machine learning. Pest Manag Sci. 2023;79: 1225–1233. doi:10.1002/ps.7296

25. Wolf SW, Ruttenberg DM, Knapp DY, Webb AE, Traniello IM, McKenzie-Smith GC, et al. NAPS: Integrating pose estimation and tag-based tracking. Methods Ecol Evol. 2023;14: 2541–2548. doi:10.1111/2041-210X.14201

26. Sunami N, Kimura H, Ito H, Hashimoto K, Sato Y, Tachibana S, et al. Automated escape system: identifying prey’s kinematic and behavioral features critical for predator evasion. J Exp Biol. 2024;227: jeb246772. doi:10.1242/jeb.246772

27. Pitera AM, Heinen VK, Sonnenberg BR, Benedict LM, Bridge ES, Pravosudov VV. Social group membership does not facilitate spatial learning of fine-scale resource locations. Behav Ecol Sociobiol. 2025;79: 104. doi:10.1007/s00265-025-03655-8

28. Lendvai ÁZ Akçay Ç, Weiss T, Haussmann MF, Moore IT, Bonier F. Low cost audiovisual playback and recording triggered by radio frequency identification using Raspberry Pi. PeerJ. 2015;3: e877. doi:10.7717/peerj.877

29. Walter T, Couzin ID. TRex, a fast multi-animal tracking system with markerless identification, and 2D estimation of posture and visual fields. Lentink D, Rutz C, Pujades S, editors. eLife. 2021;10: e64000. doi:10.7554/eLife.64000

30. Tian Y, Ye Q, Doermann D. YOLOv12: Attention-Centric Real-Time Object Detectors. arXiv; 2025. doi:10.48550/arXiv.2502.12524

31. Rosenthal R. Biasing Effects of Experimenters. ETC Rev Gen Semant. 1977;34: 253–264.

32. Burghardt GM, Bartmess-LeVasseur JN, Browning SA, Morrison KE, Stec CL, Zachau CE, et al. Perspectives – Minimizing Observer Bias in Behavioral Studies: A Review and Recommendations. Ethology. 2012;118: 511–517. doi:10.1111/j.1439-0310.2012.02040.x

33. Hróbjartsson A, Thomsen ASS, Emanuelsson F, Tendal B, Hilden J, Boutron I, et al. Observer bias in randomized clinical trials with measurement scale outcomes: a systematic review of trials with both blinded and nonblinded assessors. CMAJ. 2013;185: E201–E211. doi:10.1503/cmaj.120744

34. Rosenthal R. Experimenter effects in behavioral research, Enlarged ed. Oxford, England: Irvington; 1976. pp. xiii, 500.

35. Schulz KF, Chalmers I, Hayes RJ, Altman DG. Empirical evidence of bias: dimensions of methodological quality associated with estimates of treatment effects in controlled trials. JAMA. 1995;273: 408–412. doi:10.1001/jama.1995.03520290060030

36. Holman L, Head ML, Lanfear R, Jennions MD. Evidence of experimental bias in the life sciences: why we need blind data recording. PLOS Biol. 2015;13: e1002190. doi:10.1371/journal.pbio.1002190

37. Schulz KF, Grimes DA. Blinding in randomised trials: hiding who got what. The Lancet. 2002;359: 696–700. doi:10.1016/S0140-6736(02)07816-9

38. Kilkenny C, Parsons N, Kadyszewski E, Festing MFW, Cuthill IC, Fry D, et al. Survey of the quality of experimental design, statistical analysis and reporting of research using animals. PLOS ONE. 2009;4: e7824. doi:10.1371/journal.pone.0007824

39. Hróbjartsson A, Thomsen ASS, Emanuelsson F, Tendal B, Rasmussen JV, Hilden J, et al. Observer bias in randomized clinical trials with time-to-event outcomes: systematic review of trials with both blinded and non-blinded outcome assessors. Int J Epidemiol. 2014;43: 937–948. doi:10.1093/ije/dyt270

40. Roche DG. Effects of wave-driven water flow on the fast-start escape response of juvenile coral reef damselfishes. J Exp Biol. 2021;224: jeb234351. doi:10.1242/jeb.234351

41. Wascher CAF, Bugnyar T. Behavioral Responses to Inequity in Reward Distribution and Working Effort in Crows and Ravens. PLOS ONE. 2013;8: e56885. doi:10.1371/journal.pone.0056885

42. Marsh DM, Hanlon TJ. Observer gender and observation bias in animal behaviour research: experimental tests with red-backed salamanders. Anim Behav. 2004;68: 1425–1433. doi:10.1016/j.anbehav.2004.02.017

43. Oreg S, Bayazit M. Prone to bias: development of a bias taxonomy from an individual differences perspective. Rev Gen Psychol. 2009;13: 175–193. doi:10.1037/a0015656

44. Berman GJ, Choi DM, Bialek W, Shaevitz JW. Mapping the stereotyped behaviour of freely moving fruit flies. J R Soc Interface. 2014;11: 20140672–20140672. doi:10.1098/rsif.2014.0672

45. Khanam R, Hussain M. YOLOv11: An overview of the key architectural enhancements. arXiv; 2024. doi:10.48550/arXiv.2410.17725

46. Fujiwara T, Brotas M, Chiappe ME. Walking strides direct rapid and flexible recruitment of visual circuits for course control in Drosophila. Neuron. 2022;110: 2124–2138.e8. doi:10.1016/j.neuron.2022.04.008

47. Minasandra P, Sridhar V, Roche D, Planas-Sitjà I. TracktorLive: an integrated real-time object tracking and response system. 2026 [cited 12 Mar 2026]. Available: https://osf.io/6sxrk

