## supplementary information for "TracktorLive: an integrated real-time object tracking and response system"

### 1. Design and operation of TracktorLive

A typical pipeline using TracktorLive consists of a 'Tracktor Server' (hereafter, a server) that performs video analysis and object tracking and, optionally, a number of 'Tracktor Clients' (hereafter, clients) that carry out actions (Fig. S1). This design enables high speed parallel execution of code for object tracking (on the server side) and for handling decisions and responses (on the client side).

#### *1.1. Tracktor Server*

The server is the central part of the code (Fig. S1). It can be customised with a number of 'cassettes', which are simple python functions connected to the server through specific decorators. In Python syntax, a decorator is a small bit of text beginning with an `@` that precedes the declaration of a function. By using different decorators, the user can specify when exactly the 'cassette' function is to be executed. The server is initialised with a number of parameters related to object tracking, and upon initialisation, executes all the cassettes specified by the user to run 'at start' (connected through the decorator, `@server.startfunc`). During initialisation, users can specify a `feed ID`, a unique identifier for the server (e.g., 'fish tank 5'). If not specified, a random combination of characters is assigned as the `feed ID`.

In each iteration, when the main server process has started, a frame captured from a camera is sent to the server. First, the server executes all in-process cassettes declared to run during operation (connected through the decorator `@server`, see Fig. S2 for an example). These cassettes may be helpful for video optimisation, book-keeping, or other adjustments, such as controlling if and when data is recorded (see example 3.2 in the main text). The server then runs Tracktor [1] in the background to perform object detection and tracking, as described in the main text.

By default, the server maintains a constantly updated ‘buffer’ of frames (typically the last 10 s, but user-specifiable), tracked locations of individuals, and timestamps for each frame. The locations and timestamps are stored in shared memory protected by a semaphore. The semaphore ensures that read and write operations by the server and clients (see below) do not happen simultaneously, because such ‘race conditions’ can lead to garbled data. The semaphore must also be passed (or ‘acquired’ in programming parlance) by the server to access its own storage of previous frames. This ensures that the correct frames always correspond to the correct tracked locations and timestamps.

Upon shutdown, the server executes another cache of cassettes (connected through the decorator `@server.stopfunc`), which controls final shutdown and book-keeping operations. ‘At-start’, ‘in-process’, and ‘at-stop’ cassettes are all executed in the order in which they are declared in the script. The server can also be customised by a large number of flags (python variables ‘belonging’ to the server, accessed as, e.g., `server.write_recordings`) controlling, for instance, whether data is to be constantly written to storage, whether videos are to be saved or disposed, whether the server must halt on meeting some condition, and whether the server is initialised with a time-out beyond which it needs to stop. All flags can be specified at initialization and edited through cassettes at any time.

In one script, a user can declare multiple servers, for instance to manage a setup with multiple cameras. This can be useful in setups where multiple experimental arenas exist and need to simultaneously be recorded, or where multiple views of an experiment are needed. In such cases, the servers can be spawned with different names (e.g., server1, server2…) and then initialised individually in the script.

Additional, detailed documentation is available on the TracktorLive GitHub repository.

#### *1.2. Tracktor Clients*

Clients are far simpler than servers. A client is initialised by declaring the `feed ID` of the server to which it must connect. The client then automatically retrieves all necessary information. Clients can also be customised with 'at-start' (`@client.startfunc`), 'in-process' (`@client`, see Fig. 2 in the main text for an example), and 'at-stop' (`@client.stopfunc`) cassettes. Client cassettes are more limited, in that they cannot directly access the camera or the server. They simply get semaphore-protected access to the buffer of tracked data and timestamps exposed by the server. However, as their cassettes can contain arbitrary python code, clients can be made to perform actions in response to the tracked trajectories in the server's buffer. Since clients do not handle any video capture, video editing, or object tracking operations, their response time is quite fast. Multiple clients can be declared and connected to any server, and they will all execute their cassettes in separate parallel processes.

### 2. Experimental comparison of manual stimulus delivery versus TracktorLive

To compare tracking and stimulus delivery by TracktorLive versus human experimenters, we conducted an experiment that required participants to press a button whenever a test fish crossed the left or right boundary of an imaginary region of interest (ROI). No stimulus was delivered to the test fish but we recorded and compared the timing of the activation of the fictitious stimulus by experimenters to that of the Arduino board controlled by TracktorLive. Two fish species were tested: medaka (*Oryzias latipes*), which exhibit slow and predictable movements, and neon tetra (*Paracheirodon innesi*), which frequently engage in burst swimming. The fish were sourced from an office aquarium in the laboratory of Animal Ecology (department of Biological Sciences, Tokyo Metropolitan University), and had been purchased from a local supplier.

The experimental arena consisted of a white, opaque aquarium (30 × 20.5 × 5 cm) filled with 2.5 cm of water. The arena was placed inside a wooden box (60 × 25 × 29 cm) to minimize external stimuli. A webcam (EMEET HD1080P) was affixed 35 cm above the arena via a Manfrotto articulated arm with a camera bracket, and connected to a laptop on an experimental bench.

A single fish was introduced into the arena and left to habituate for five minutes while we initialised the tracking software and briefed a human experimenter on the task. The experimenters (three men, three women) were undergraduate and graduate students from the department of Biological Sciences of the Tokyo Metropolitan University. Experimenters monitored the test fish's movements and location via a live feed to the laptop.

The experimental arena was marked with black lines at 9 cm and 22.5 cm along its length (30 cm), indicating the boundaries of the region of interest (ROI), at the extremities of the arena (Fig. S3A). When the experimenter pressed the `m` key on the keyboard, a red light—visible to the webcam—was activated. This allowed us to assess response accuracy by measuring the time difference between the fish crossing a line delimiting the boundaries of the ROI and the activation of the light by the experimenter. Each experimenter completed a five-minute trial during which the test fish could cross the lines repeatedly (the number of crossings per trial can be seen in Table S1). Three experimenters conducted the trial using medaka fish, while the other three experimenters used tetra fish. To minimise bias among participants, TracktorLive processed the webcam feed in real-time, displaying a red dot on the fish's centre of mass.

After all three experimenters conducted their trial with medaka fish, we ran a five-minute trial with the same fish using TracktorLive, programming it to trigger the signal when the fish was detected within a ±10 px range of the ROI boundaries. We followed the same procedure with tetra fish. The performance of both experimenters and TracktorLive was manually evaluated by a researcher (blind to the treatment) using a custom Python script (available in our Open Science Framework repository [2]). Briefly, the researcher paused the video at the frame when the red light turned on, clicked on the dot at the centre of mass of the fish, and resumed the video. During this process, the script automatically recorded the pixel position of the fish's centroid (previously clicked by the researcher) and calculated its distance from the actual boundary (which remained invisible to the evaluator).

We used a generalised linear mixed-effects model with a negative binomial error distribution [3] to test whether the method (experimenter or TracktorLive) and fish species affected signal-triggering accuracy. We used error distance (in pixels) as the response variable, and specified method and fish species as fixed effects. Experimenter identity was included as a random factor with six levels. Model assumptions and overdispersion were checked with model diagnostics generated by the package [4]. Finally, we used two Kruskal–Wallis tests to assess whether experimenters differed significantly in accuracy, one for each fish species.

Both human experimenters and TracktorLive successfully triggered the fictitious stimulus at the correct boundaries, with mean trigger positions close to the true boundaries of the ROI (Fig. S3A). However, TracktorLive demonstrated consistently higher accuracy and lower variability compared to human experimenters, particularly with the medaka fish, which moved more slowly (Fig. S3A,B).

The overall error distribution, defined as the distance between the triggered pixel and the true boundary, was about twice narrower, and four times lower (Table S1, Fig. S3A,B), for TracktorLive than for human experimenters for both medaka and tetra fish. Experimenters exhibited significantly higher error rates than TracktorLive (GLMM: $\beta = 1.06 \pm 0.45, z = 2.34, p = 0.02$), and accuracy was significantly lower in tetra fish than medaka fish ($\beta = 0.73 \pm 0.20, z = 3.63, p < 0.001$), with no evidence for an interaction between fish species and method. Furthermore, human responses exhibited substantial inter-individual variation for both medaka (Kruskal-Wallis: $\chi^2_{2,39} = 7.22, p = 0.02$) and tetra ($\chi^2_{2,64} = 242.16, p < 0.001$) fish trials (Fig. S3B).

These results highlight the advantages of using TracktorLive over manual observation and stimulus operation: the greater variability observed in human responses suggests that experimenter-dependent differences could influence behavioural outcomes, potentially affecting the reliability of experimental conclusions.

### References

1. Sridhar VH, Roche DG, Gingins S. Tracktor: Image-based automated tracking of animal movement and behaviour. Methods in Ecology and Evolution. 2019;10: 815–820. doi:10.1111/2041-210X.13166

2. Minasandra P, Sridhar V, Roche D, Planas-Sitjà I. TracktorLive: an integrated real-time object tracking and response system. 2026 [cited 12 Mar 2026]. Available: https://osf.io/6sxrk

3. McGillycuddy M, Popovic G, Bolker BM, Warton DI. Parsimoniously Fitting Large Multivariate Random Effects in glmmTMB. Journal of Statistical Software. 2025;112: 1–19. doi:10.18637/jss.v112.i01

4. Hartig F. DHARMa: Residual Diagnostics for Hierarchical (Multi-Level / Mixed) Regression Models. 2026. Available: https://github.com/florianhartig/DHARMa

**Table S1: Quantitative comparison of error in stimulus triggers between TracktorLive and human participants.** Comparison of the error, measured as the distance (in px) between the pixel at which the stimulus was triggered and the pixel representing the true boundary, for experimenters (manual) and TracktorLive (automated) triggering the fictitious stimulus using medaka and tetra fish.

| | **Mean error (px)** | **Standard deviation of error (px)** | **Number of observations** |
|---|---|---|---|
| **Medaka fish** | | | |
| Experimenters | 9.9 | 8.41 | 42 |
| TracktorLive | 2.59 | 2.08 | 22 |
| **Tetra fish** | | | |
| Experimenters | 14.46 | 12.07 | 67 |
| TracktorLive | 5.9 | 4.09 | 40 |

**Table S2: Potential use cases for TracktorLive.**

| Area | Application / Use-case | Cassette / Functionality Needed |
|---|---|---|
| **Behavioural experiments** | Escape-response experiments | - Tracking-based stimulus delivery cassettes |
| | Animal cognition & choice / decision-making experiments (adapt environment depending on animal choice or action) | - Cassette to modify environmental variable (light, temperature, etc) |
| | Virtual-reality setups for single individuals | - Experiment-design cassette that synchronises VR output with behavioural tracking |
| | Detect sick / dead lab animals | - Health-monitoring cassette (motion / behaviour analysis) |
| | Automatic feeders / enrichment devices | - Tracking & Arduino control cassettes, enrichment-control cassette (periodically drives feeders, toys, etc.) |
| **Experimenter ease-of-life** | Transmit tracking data over a network port | - Network-streaming cassette (TCP / UDP broadcast) |
| | Email notifications when events of interest occur | - Email-notification cassette |
| | Daily visualisations of animal movement | - Daily-summary visualisation cassette |
| **Extensions of TracktorLive** | Smart traps (triggered only when the target species is recognized) | Species-recognition cassette using deep learning tools (lightweight classifier) |
| | Welfare monitoring in animal facilities (detect repetitive stereotypic movement) | Movement-pattern cassette (detects repetitive trajectories, notifies management) |

| | | |
|---|---|---|
| | Fish-welfare in aquariums (adjust oxygenation / water replacement based on overall motion) | - Movement-pattern cassette (overall activity detection)<br>- Environmental-control cassette (modulates oxygenation, starts water exchange) |
| | Centring in optical devices (Arduino-based stage / telescope dial control) | Motor-control cassette (drives stepper / servo motors) or video centring cassette.<br>- Vision-feedback cassette (continuous centroid tracking to keep target centred) |
| | Baby wake/sleep monitor (privacy-preserving alert + soothing sounds/white-noise when baby exceeds motion threshold or crib rocking activation) | - Motion-threshold or sound-recognition cassette<br>- Alert-action cassette (push notification)<br>- Action cassette (audio playback or motor activation to start gentle rocking of the crib) |

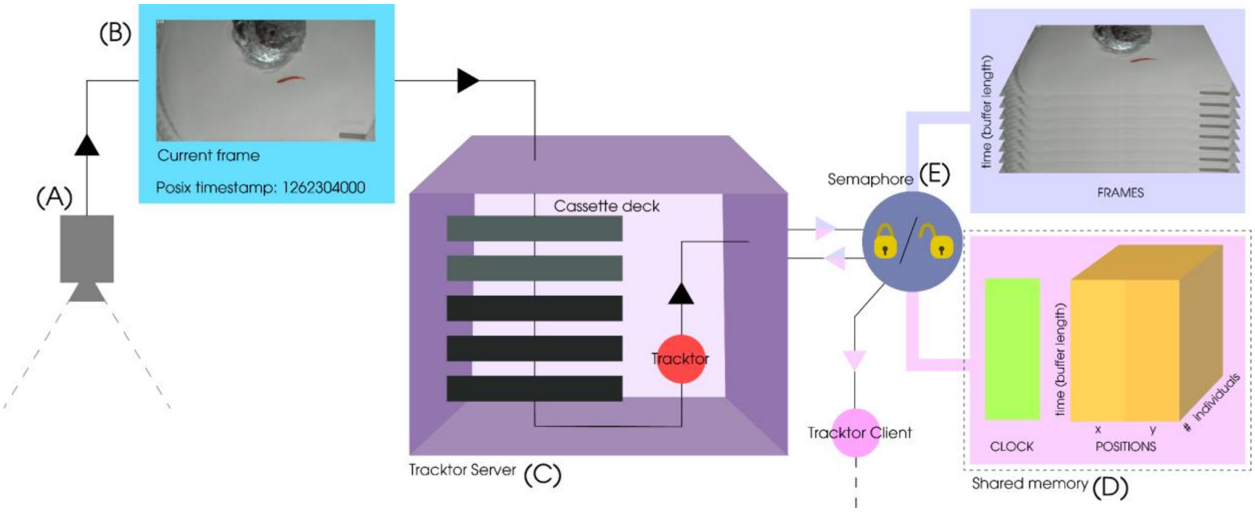


**Figure S1: The software architecture of TracktorLive.** In a typical workflow, a camera **(A)** captures a video frame **(B)** and transmits it to the pre-initialised Tracktor server **(C)**. The server then applies specified cassettes that could, for instance, edit the incoming frame, or make recordings based on other sensors. The server then internally uses Tracktor [1] to track one or multiple objects and stores their tracked location(s) as well as the timestamp of the video frame in a shared memory **(D)**. The server also stores the video frame itself. The last 10 s (or another user-specified interval) of data (CLOCK, POSITIONS) and frames (FRAMES) are stored, deleting the oldest items at every iteration. To write or read data from the shared memory, the server must pass a semaphore 'lock' **(E)**, which ensures that data retrieved is synchronous (the indices of the frames correspond directly to the indices of the tracked locations and timestamps), and not accessed by multiple parallel processes simultaneously.

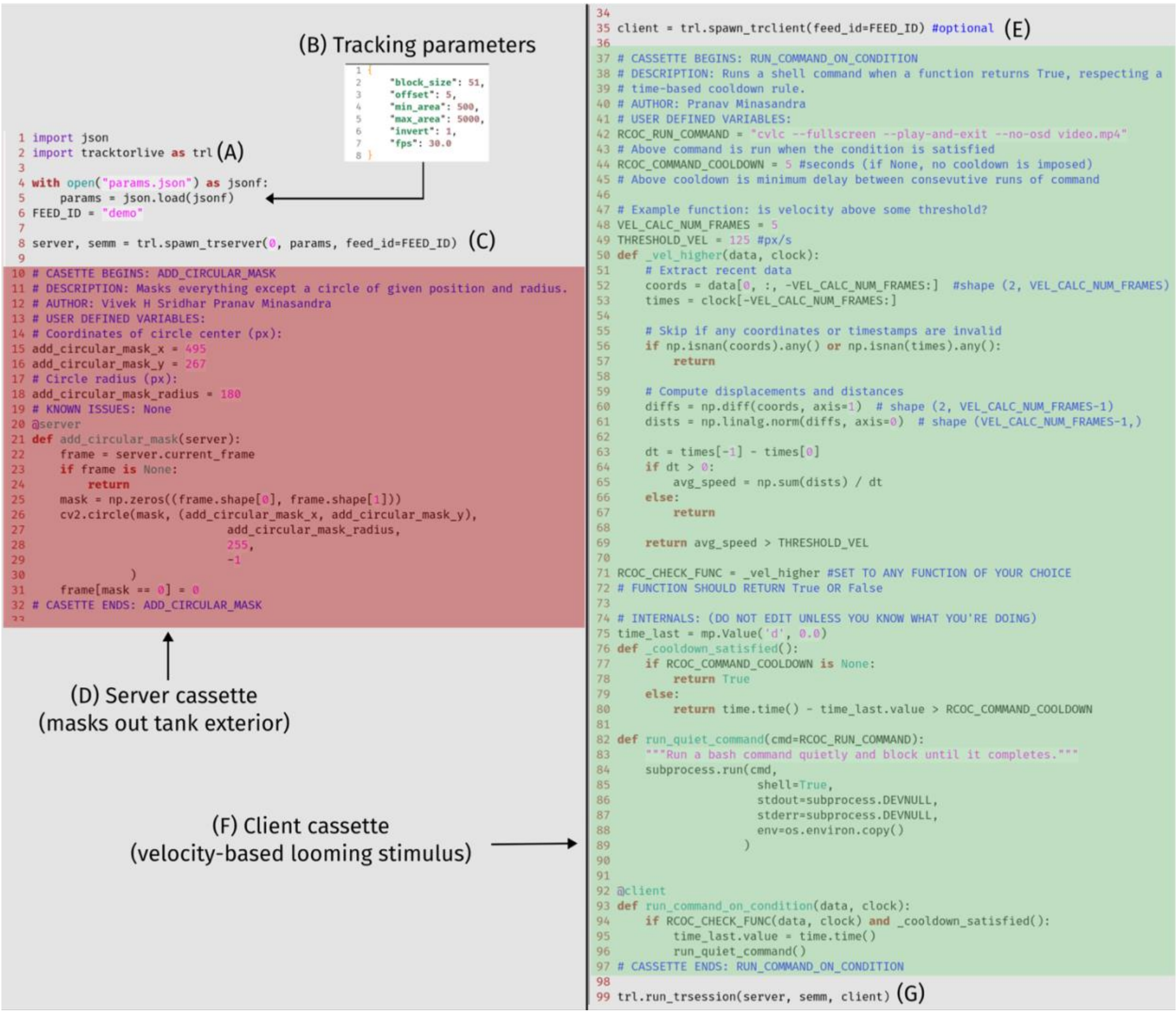


**Figure S2: A typical python script for using TracktorLive.** This example script is used to optimally track a fish in a tank and plays a video whenever the fish's velocity exceeds a certain threshold. **(A)** Tracktorlive is imported as `trl` by convention. **(B)** Tracking-related parameters are read in, and **(C)** passed to a function that spawns a Tracktor Server and a semaphore manager. **(D)** Server cassettes (here, a circular mask) are attached to the server. Users can either write their own cassettes, or simply choose, copy, and paste cassette code from our GitHub repository's Library of Cassettes. At this point, tracking has already been set up. **(E)** Clients can now be added to adjudicate response delivery. **(F)** Client cassettes, which can also be written by users or chosen from the online Library of Cassettes, are added (here, the velocity-based presentation of a looming stimulus). **(G)** Finally, both the server and the client spawned before are initialised.

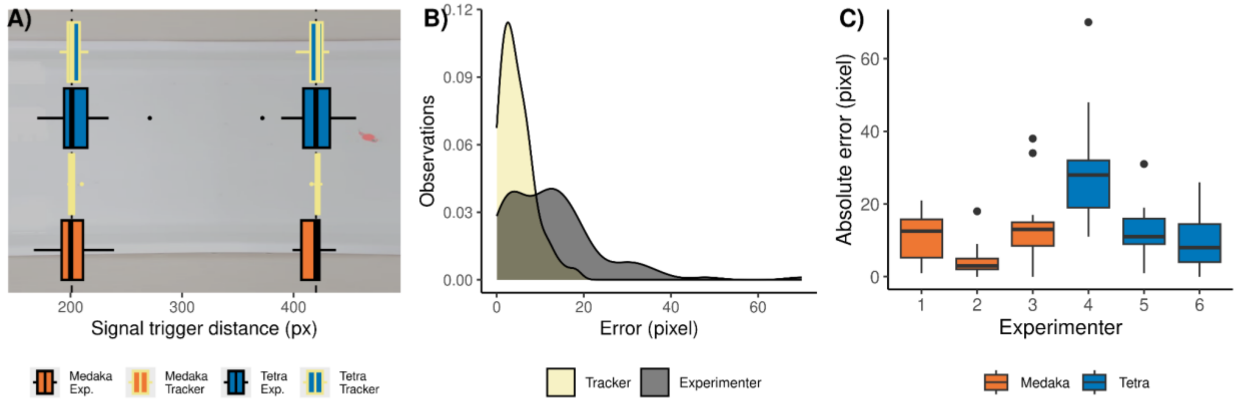


**Figure S3: Comparison of the accuracy in the timing of stimulus delivery by human experimenters and automated tracking with TracktorLive. (A)** Black vertical dotted lines are the left and right boundaries of the region of interest. The stimulus had to be triggered when the fish's centre of mass crossed either line. Boxplots indicate the median and inter-quartile range. **(B)** The distribution of the absolute error in pixels (observed – expected pixel) for manual (grey) and automated (yellow) stimulus activation. **(C)** Absolute error distribution for manual stimulation of medaka (orange) and tetra (blue) fish by six experimenters. The number of observations for the trials with human experimenters and automated tracking are indicated in Table S1.
